# Patterning of membrane adhesion under hydraulic stress

**DOI:** 10.1101/2023.01.04.522479

**Authors:** Céline Dinet, Alejandro Torres-Sánchez, Marino Arroyo, Margarita Staykova

## Abstract

Hydraulic fracturing plays a major role in the formation of biological lumens during embryonic development, when the accumulation of pressurized fluid leads to the formation of microlumens that fracture cell-cell contacts and later evolve to form a single large lumen. However, the physical principles underpinning the formation of a pattern of microlumens from a pristine adhesion and their subsequent coarsening are poorly understood. Here, we use giant unilamellar vesicles adhered to a supported lipid bilayer and subjected to osmotic stress to generate and follow the dynamics of hydraulic fracturing akin to those in cells. Using this simplified system together with theoretical modelling and numerical simulations, we provide a mechanistic understanding of the nucleation of hydraulic cracks, their spatial patterns and their coarsening dynamics. Besides coarsening, we show that microlumens can irreversibly bud out of the membrane, reminiscent of endocytic vesicles in cell-cell adhesion. By establishing the physics of patterning and dynamics of hydraulic cracks, our work unveils the mechanical constraints for the biological regulation of hydraulically-driven adhesion remodeling.

## Introduction

Hydraulic pressure is a major force at cellular and tissue scales^1;2;3^ compromising tissue integrity^4^ and driving cell blebbing^5^, cell fate decisions^6;7^, embryo development^6;8;9^, or organ morphogenesis^10;11;12^. Some of these processes rely on the ability of hydraulic pressure to selectively detach cell-cell or cell-matrix adhesions. For instance, hydraulic cell-cell fracture is thought to determine the first stages of mammal development, during which pressurization of the gaps at cell-cell junctions produce a widespread distribution of small pockets, which subsequently undergo an actively guided coarsening process to form the blastocoel^8^. Immediately subsequent stages of development involve further luminogenesis and water management between lumens^9^. In-vitro studies have shown that cell-autonomous^13;14^, poroelastic^4;15^, or osmotically applied^15;16;17^ pressure differences can lead to patterns of pressurized pockets of various sizes, from sub-micron cracks to multicellular cavities.

The nature of the pattern of hydraulically driven pockets should play a major role in determining the subsequent shape of organs (network of bile ducts), the resilience of the epithelial barrier under hydraulic stress or stretch^4;18^ or the robustness of morphogenesis^8^. The formation of an initial pattern of hydraulic cracks should be a largely physical process, which then cells and tissues may be able to control by biologically tuning in space and time physical parameters^19^. The physics of coarsening of an array of preexisting water pockets has been studied^8;20^. However, the physical principles controlling when and how a pattern of hydraulic cracks emerges in the first place from a pristine adhesion remain largely unknown.

To identify these principles, we combine experimental observations on adhered lipid vesicles, with mathematical and computational modelling. The hydraulic fracture in embryonic tissues is driven by pressure gradients established through active ion transport across cell membranes, followed by a passive compensatory efflux of water into the cell-cell interstice^8^. To generate such pressure gradients in our artificial system, we subject the vesicles to osmotic shocks and observe hydraulic fracturing dynamics akin to those in cells. We present a comprehensive picture of the various mechanical and transport mechanisms controlling the formation of membrane hydraulic cracks, their spatial patterns and coarsening dynamics and show how they can be controlled by the magnitude of pressure gradients, and the type and density of membrane bonds.

## Results

### Hydraulic fracturing in lipid vesicles

In our default experimental setup, we study the hydraulic fracturing between a giant lipid vesicle (GUV) adhered to a supported lipid bilayer (SLB). Both the vesicle and the SLB, contain biotinylated lipids (b) at a desired density (0.2, 1 or 4 mol%). Once the SLBs are deposited on the glass substrate, they are exposed to fluorescent Neutravidin-DyLight 488 (NAV), which bind the biotin groups (b-NAV bond). Excess NAV is then washed from the medium. When the vesicles are added on top of the SLB they adhere to it by forming b-NAV-b bonds (Fig. 1a,b). Most bonds appear mobile, as confirmed by FRAP experiments, see Supplementary Information (SI). Consistent with previous studies on enrichment of mobile bonds in the adhesion zone^21;22^, we observe that the NAV fluorescence intensity between the two membranes is between 1-2.5 times higher compared to that on the bare SLB (Fig. 1a and SI).

**Figure 1:**
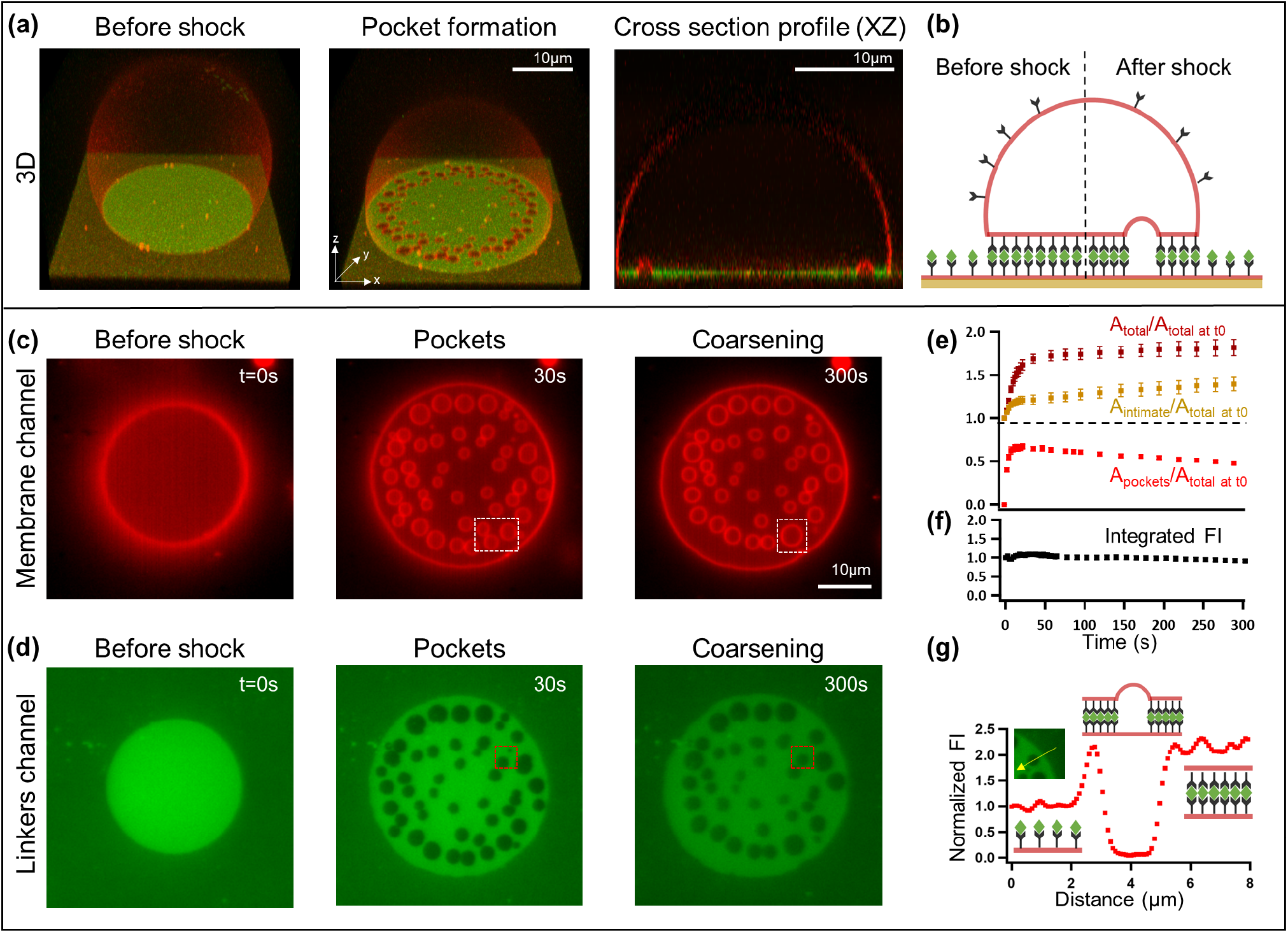
Hydraulic fracturing of adhered lipid membranes. (a) 3D images of a vesicle bound to a supported membrane (at 1 mol% biotinylated lipids) before and after the application of a hyper-osmotic shock of 100 mM; cross-section of the vesicle after the shock. The membranes, labelled by Rhodamine, appear in red and the NAV bonds, labelled by DyLight488, in green. (b) Sketch of the vesicle and the adhesion complexes, before and after the shock. The GUV and the SLB membranes are represented in red with the biotinylated lipids in black. Neutravidin linkers are shown as green rhombuses. (c-d) Images of the membrane (c) and the corresponding NAV distribution (d) in the adhesion zone at 0 (immediately before the shock), 30 and 300 sec after the shock. The dashed squares show pocket fusion (white) and pocket collapse (red) events. (e) Total adhesion area, total fractured pocket area, and intimate adhesion area over time. The intimate area is defined as total adhesion area minus pocket area. (f) Integrated fluorescent density of NAV in the adhesion zone over time. (g) Fluorescent intensity profile of NAV across a pocket along the yellow arrow in the inset.

Vesicles that are stably adhered to the SLBs have the shape of truncated spheres (Fig. 1a) with a contact angle of ∼ 120° and a diameter of the adhesion zone of ∼ 20 μm (Fig. 1a). Upon hyper-osmotic shock, applied by rapidly increasing the concentration of osmolytes in the external vesicle medium, we observe the formation of multiple fluid-filled pockets between the two membranes, which in cross-section resemble spherical caps protruding into the vesicle (Fig. 1a). The pockets vary in size and have a non-homogeneous distribution (Figure 1c, Video 1). In the NAV fluorescent channel the pockets’ footprints appear dark (Fig. 1d, Video 1). The total fractured area, quantified by measuring the combined footprint area of the pockets, reaches its maximum in few tens of seconds after the osmotic shock (Fig. 1e). Thereafter it gradually decreases as a result of coarsening of the pocket pattern through either pocket coalescence or pocket collapse. The latter has been interpreted as a process akin to Ostwald ripening during which the collapsing pocket transfers its content to neighboring pockets by diffusive water transport in the tightly adhered interstice^8^ (Fig. 1c,d, Video 1). The total vesicle adhesion area increases simultaneously with the growth of the pockets, reaching a plateau at later times. At the same time, the intimate adhesion area of b-NAV-b membrane contacts also increases in the first tens of seconds but by a much lesser amount, followed by more gradual increase as pockets coarsen (Fig. 1e). We conclude therefore that upon hydraulic fracturing, the excess of vesicle membrane area is recruited mostly in the growth of pockets, and not in a homogeneous growth of vesicle contact area, as previously reported^23^. Similar vesicle fracturing is consistently observed for various other osmotic shocks and bond concentrations (SI). Passivation of the membranes using PEG-ylated lipids to avoid non-specific adhesion^22;24^ does not affect the formation of patterns of hydraulic cracks either (SI).

Next we discuss the dynamic behaviour of bonds during hydraulic fracturing. The symmetric shapes of the membrane pockets in our system and their wide footprint areas, remarkably resemble the microlumens observed between embryonic cells^8^. Hydraulic fractures of various shapes have also been reported for cells and vesicles adhered through immobile bonds^15;16;25^ or via non-specific adhesion^26^. To establish the role of lateral mobility of bonds in our system, we examine the NAV signal as a measure of bond density, which drops to zero at the location of the pockets from the moment of their opening (Fig. 1g) until the later stages. This suggests that as the membranes peel away and form pockets, the advancing contact line of the pocket pushes away the strong, mobile biotin-NAV bonds, thus leading to a large pocket footprint. During this redistribution, bonds are not broken and the overall number of bonds in the adhesion zone stay the same as confirmed by the constant integrated NAV density measured over the whole patch area (Fig. 1f). Furthermore, the nearly uniform NAV fluorescence intensity during pocket formation suggests fast lateral equilibration of bonds relative to the dynamics of pocket formation and evolution.

### Theoretical model

To understand pattern formation and dynamics, we develop a mathematical model predicting the nucleation and evolution of hydraulic pockets starting from a pristine adhesion patch and following an osmotic perturbation. We first examine the adhesion strength by estimating the nondimensional number 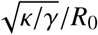 where κ is the bending rigidity, *γ* the adhesion energy and *R*_0_ the typical vesicle radius^27;28^. For adhesion mediated by mobile bonds, the effective adhesion energy is the osmotic 2D pressure of bonds trapped within the adhesion patch, approximated by *γ* = *k*_*B*_*Tc* where *k*_*B*_ is Boltzmann’s constant, *T* absolute temperature and *c* the number density of bonds^29;30^. In our system, 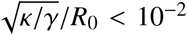 (SI) and hence vesicles are in a strong adhesion regime^27;28^, in agreement with the sharp contact angles and absence of noticeable fluctuations. We thus neglect membrane bending rigidity and fluctuations, and model the unattached part of the vesicle as a spherical cap of radius *R* and contact angle *θ* (Fig. 2a).

**Figure 2:**
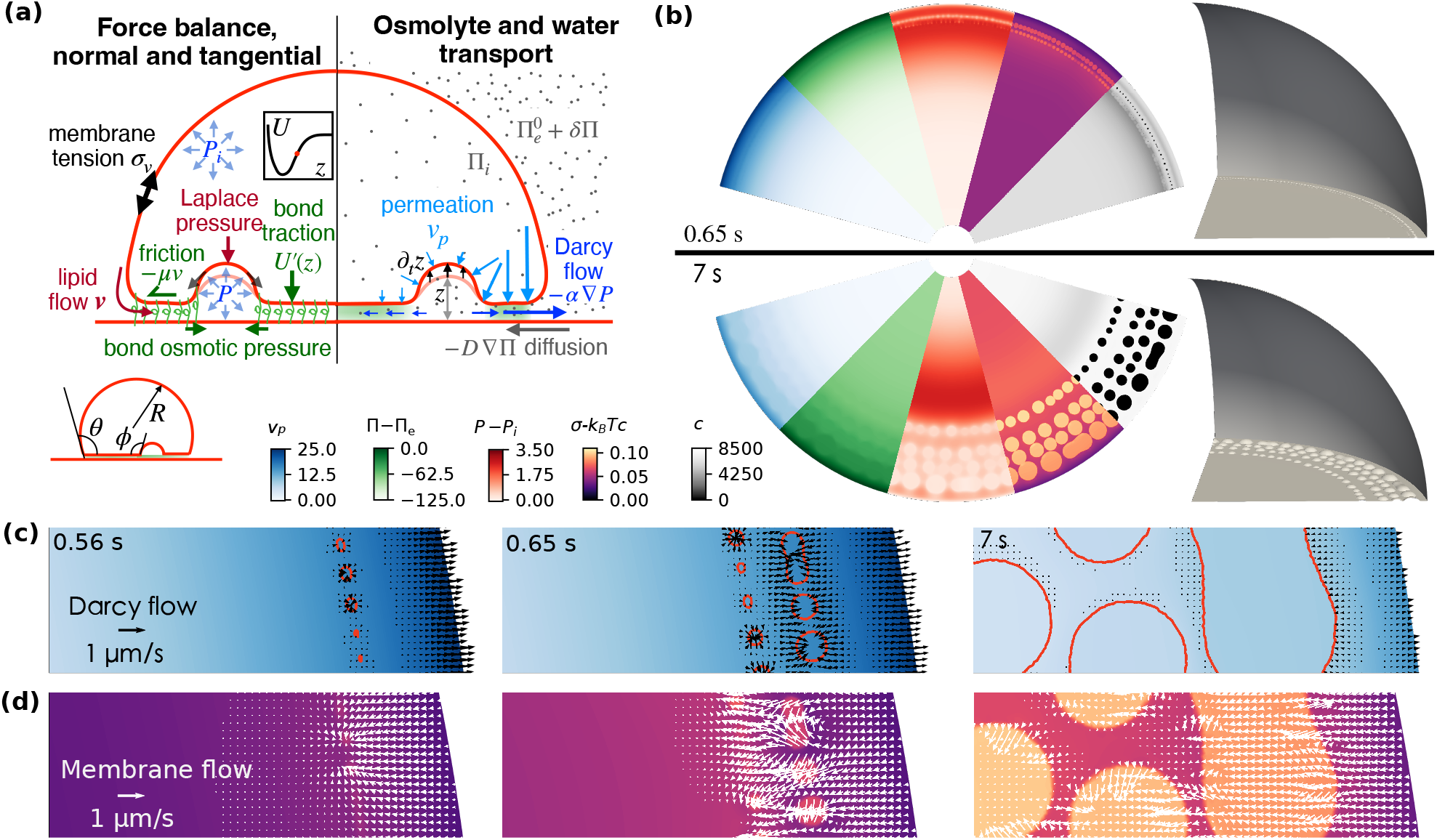
Mechanisms of hydraulic fracturing. (a) Schematic of the physical ingredients controlling the formation of a pattern of hydraulic cracks, Box 1. Mechanical ingredients are illustrated on the left and transport ingredients on the right. (b) Snapshots of various fields at the adhesion patch and 3D view of the membrane. The blue map is permeation velocity (nm s^−1^), proportional to the water chemical potential across the membrane, *P*(*x, y, t*) − Π(*x, y, t*) − (*P*_*i*_ − Π_*i*_). The green map is the osmotic pressure (kPa) relative to that of the external medium Π(*x, y, t*) Π_*e*_. The red map is the hydraulic pressure (kPa) relative to the vesicle pressure, *P*(*x, y, t*) − *P*_*i*_, whose gradient is proportional to Darcy flow. The purple map is the net membrane tension *σ* − *k*_*B*_*Tc* (kPa μm), and the gray map is the bond concentration (μm^−2^). (c) Zoom of water transport in the interstice, with water permeation (map) and Darcy flow (black arrows); pockets are outlined in red color. (d) Zoom of membrane tension (map) and membrane flow (white arrows).

Before the osmotic shock, the vesicle, the solvent, the osmolytes and the bonds are in thermodynamical equilibrium^29;30;33^ (Fig. 2a and SI). Briefly, osmolytes cannot cross the semi-permeable membrane and hence sustain an osmotic pressure difference ΔΠ = Π_*i*_ − Π_*e*_ between that of the external medium Π_*e*_ and that of the interior of the vesicle Π_*i*_ = *k*_*B*_*TN*_*o*_/*V* with *N*_*o*_ the number of trapped osmolytes and *V* the vesicle volume. Instead, water can equilibrate across the membrane, and hence its chemical potential inside, which is proportional to *P*_*i*_ − Π_*i*_, is equal to that outside, leading to ΔΠ = Δ*P*. The hydraulic pressure difference is resisted by membrane tension following Laplace’s law 2*σ*_*v*_/*R* = Δ*P*. The edge of the adhesion patch is also a closed semi-permeable interface, albeit of lower dimensionality, which allows lipids to cross but traps bonds, whose number *N*_*b*_ is assumed to be fixed (Fig. 1f,g), and which exert an osmotic tension on the edge resisted by membrane tension following a Young-Dupré relation *k*_*B*_*Tc* = *σ*_*v*_(1 − cos *θ*), where *c* = *N*_*b*_/*S* and *S* = *πR*^2^ cos^2^ *θ* is the area of the patch.

Right after the osmotic shock, Π_*e*_ suddenly increases by δΠ, bringing the system out of equilibrum. The osmotic imbalance drives water efflux through the membrane, changing the vesicle shape (*V, R, θ, S*), and hence modifying hydraulic and osmotic pressures in the vesicle, osmotic tension in the patch, and membrane tension (Π_*i*_, *P*_*i*_, *k*_*B*_*Tc* and *σ*_*v*_) according to the principles outlined above. Because the excess osmolytes from the shock can penetrate the interstice, water can also drain from the vesicle into the cleft and pressurize this space.

#### Box 1

Theoretical model

##### The adhesion patch

on the SLB is a disk 𝒟(*t*) in the (*x, y*) plane of radius *r*(*t*) = *R*(*t*) cos *θ*(*t*). The adhesion patch on the neighboring vesicle Γ(*t*) is described by the height function *z*(*x, y, t*) with (*x, y*) ∈ 𝒟(*t*). Local areas on Γ(*t*) and 𝒟(*t*) are related by *dS* = *j dxdy* where 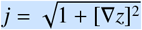. Here ∇ = (∂_*x*_, ∂_*y*_), whereas the surface nabla operator is ∇_*S*_.

##### Vesicle unknowns

Radius of spherical cap *R*(*t*), contact angle *θ*(*t*), tension *σ*_*v*_(*t*), osmotic pressure Π_*i*_(*t*) and hydraulic pressure relative to that in the external medium *P*_*i*_(*t*).

##### Unknowns in the adhesion patch

membrane height *z*(*x, y, t*); osmotic pressure Π(*x, y, t*) and relative hydraulic pressure *P*(*x, y, t*) in the interstice; tangential velocity ***v***(*x, y, t*), bare tension *σ*(*x, y, t*) and bond number density *c*(*x, y, t*) on the membrane Γ(*t*).

##### Bond distribution in an adiabatic approximation

Assuming bonds redistribute fast compared to osmolytes, water and membrane, and accounting for conservation of their number *N*_*b*_, the equilibrium Boltzmann distribution is given by

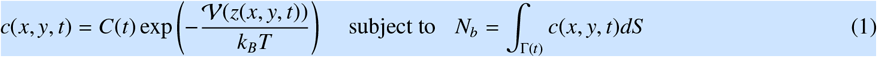

where 𝒱(*z*) is the stretching potential of a bond and the second equation determines the normalization constant *C*(*t*).

##### Mass conservation of water

According to conservation of incompressible water

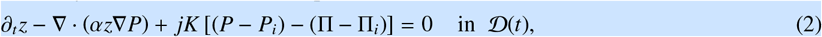

height changes must be balanced by lateral water flow following Darcy’s law ***v***^fluid^ = − *α*∇*P* with mobility *α* (second term) and by water permeation across Γ(*t*) with permeability *K* (third term).

##### Mass conservation of osmolytes

Assuming a simple van’t Hoff relation, fast equilibration along *z*, integrating through the thickness and accounting for diffusion and advection with fluid velocity following Darcy’s law, it reads

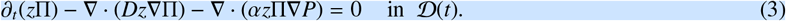

##### Tangential force balance

accounting for variations in the 2D membrane stress and for friction reads

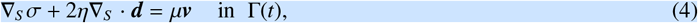

where η is the membrane viscosity, ***d*** the rate-of-deformation tensor and μ the friction coefficient. For a deforming surface, ***d*** = (_*S*_ ***v*** + (_*S*_ ***v***)^*T*^)/2 − *v*_*n*_ ***k***, where *v*_*n*_ = (∂_*t*_*z*)/ *j* is the normal velocity and ***k*** the curvature tensor^31;32^. In our convention, the normal vector to Γ(*t*) points into the vesicle and curvature of a pocket is negative. *σ* is the Lagrange multiplier field enforcing membrane inextensibility 0 = ∇_*S*_· ***v*** − *v*_*n*_*H* where *H* is the mean curvature.

##### Normal force balance

accounting for hydraulic and Laplace pressures and for bond traction reads

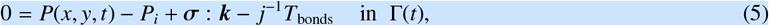

where the full 2D stress tensor supported by the membrane is *σ* = (*σ* − *γ*) ***g*** + 2η***d*** with *γ* = *k*_*B*_*Tc* the 2D osmotic pressure of bonds, and the bond traction on Γ(*t*) along *z* is *T*_bonds_ = *c*𝒱′(*z*).

##### Vesicle-scale mechanics and mass conservation

Mechanical force balance in the free-standing part is given by Laplace’s law. Force balance at the edge of the patch is given by a Young-Dupré-like equation 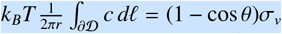, where the right-hand-side is the average 2D osmotic pressure of bonds along the edge. Conservation of incompressible enclosed water imposes that the rate of change of volume of the vesicle is balanced by permeation in the free-standing spherical cap and in the adhesion patch, and conservation of inextensible lipids imposes that the total vesicle area remains constant. Finally, conservation of the number of trapped osmolytes *N*_*o*_ imposes that Π_*i*_*V*_*i*_ = (*k*_*B*_*T*)*N*_*o*_ is constant.

##### Boundary conditions at the edge of the patch

Because the edge of the patch is not an obstacle for water or osmolyte transport between the external and interstitial media, continuity of hydraulic and osmotic pressures provides boundary data, *P*|_∂D_ = 0 (external hydraulic pressure is reference) and Π|_∂D_ = Π_*e*_. Similarly, lipids can flow through the edge, and hence *σ*|_∂D_ = *σ*_*v*_.

##### Initial conditions

Starting from an equilibrium state for 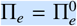 for all unknowns, we suddely increase external osmotic pressure to 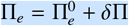 at *t* = 0 and self-consistently solve all the equations above over time.

##### Model parameters

Mass: number of bonds *N*_*b*_, number of trapped osmolytes *N*_*o*_, and vesicle surface area. Osmotic pressures: 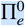 and shock magnitude δΠ. Membrane properties: viscosity η and permeability *K*. Bonds: stretching potential 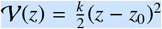 with *k* the stiffness and *z*_0_ the resting separation. Interstice: Darcy mobility *α*, diffusivity *D* and friction μ. To account for the fact that in detached regions, where *c* ∼ 0, bare membrane tension and hydraulic pressure should equilibrate instantly, and hence friction and inverse Darcy mobility should vanish, we assume the relations μ(*c*) = μ_0_*c*/*c*_0_ and *α*(*c*) = *α*_0_*c*_0_/*c*, where μ_0_ and *α*_0_ are reference values at the nominal concentration *c*_0_.

To account for the physics of the adhesion patch, we develop a detailed model allowing us to determine the osmotic and hydraulic pressures in the interstitial space, the bond density, and vesicle membrane mechanics (its shape, tension and lipid flow). This model self-consistently couples transport and mechanical phenomena in the adhesion patch and in the free-standing vesicle (Fig. 2a, Box 1 and SI). Briefly, we align the (*x, y*) plane with that of the SLB and assume fast equilibration of osmotic and hydraulic pressures in the interstice along the thin *z* direction. Transport of osmolytes is controlled by diffusion and advection of the aqueous solution, Eq. (3). To model water transport, we view the thin and crowded interstice as a porous medium where water moves against gradients of hydraulic pressure according to Darcy’s law^34^. Then, balance of mass of the incompressible solution expresses the fact that changes in membrane height should be balanced by lateral Darcy flow and by water permeation across the membrane, Eq. (2). The fluid membrane supports a tangential stress tensor including bare tension, the osmotic tension of bonds, and viscous stresses. Variations in membrane stress generate tangential forces balanced by friction, Eq. (4), which physically results from the resistance to membrane flow posed by an array of obstacles, here the bonds^35^. Membrane stress also generates a Laplace pressure normal to the membrane, balanced by the difference of hydraulic pressures across the membrane and by the traction due to bond stretching *T*_bonds_ keeping the membranes together, Eq. (5). Consistent with our observations, we assume bond distribution adiabatically equilibrates reaching a Boltzmann distribution, Eq. (1).

### Mechanisms of hydraulic fracturing

To theoretically understand the formation of patterns of hydraulic cracks, we discard linear stability analysis^17^ given the lack of uniformity and the large perturbations in our system. Instead, we develop a finite element method for the model above in its full nonlinearity without additional assumptions (SI). In a reference simulation, we used parameter values consistent with our experiments and the literature, Supplementary Table 1 and SI.

Starting from an equilibrated adhesive vesicle, our numerical simulations readily develop arrays of hydraulic cracks upon osmotic shock application, which closely resemble our experimental observations, Fig. 2b,c and Video 2. Our simulations give us access to all physical fields with high temporal and spatial resolutions, thus allowing us to examine in detail the initial stages of pocket formation. Right after the shock, diffusion from the outer medium rapidly increases the osmotic pressure in the margin of the adhesion patch (green map in Fig. 2b). Without time to change *z*(*x, y, t*) significantly, the last two terms of Eq. (5) remain unchanged and hence so does *P*(*x, y, t*) − *P*_*i*_. As a result, the local increase of Π(*x, y, t*) is mirrored by a decrease of water chemical potential across the membrane (*P* − Π − (*P*_*i*_ − Π_*i*_)) in the margin, which drives permeation efflux into the interstitial region (blue map in Fig. 2b,c). Very close to the edge, water can escape to the outer medium (black arrows in Fig. 2c). However, at a distance from it, water is hydraulically confined by Darcy resistance and accumulates by increasing locally *z*.

The local swelling of the interstitial space stretches bonds, increases the traction they support, *T*_bonds_ = *c*𝒱′(*z*) where 𝒱(*z*) is the stretching potential of an individual bond, and pressurizes the interstice. However, increasing separation also decreases bond concentration according to Eq. (1) (gray map in Fig. 2b), which reduces the ability of bonds to resist pressure. This effect can be understood by noting that bond traction *T*_bonds_(*z*) = *U*′(*z*) derives from an effective potential *U*(*z*) = −*k*_*B*_*TC* exp [−𝒱(*z*)/(*k*_*B*_*T*)], which even if bonds visit a nearly quadratic potential with stiffness *k*, is non-convex and can sustain a maximum threshold pressure 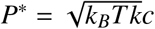 at critical separation 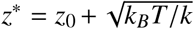. Beyond this point, the bond ensemble looses adhesion stability. Thus, from the interplay between permeation and hydraulic confinement, this condition is met at a distance from the edge, leading to the nucleation of fractured regions devoid of bonds surrounded by a region of intimate adhesion rich in bonds (gray map in Fig. 2b). The theoretical system is initially axisymmetric and remains axisymmetric up to the onset of fracture with a ring-like peeling zone. This ring, however, rapidly splits into droplet-like spherical pockets as a result of a symmetry-breaking transition akin to a Rayleigh-Plateau instability, Video 2.

The formation of pockets locally relaxes hydraulic pressure (red map in Fig. 2b), which drives Darcy water flow towards pockets to sustain their growth (black arrows in Fig. 2c and Video 3). Bonds being pushed away from detached regions, the remaining hydraulic pressure can only be resisted there by bare membrane tension through the Laplace term, Eq. (5), which recruits membrane area by lipid flows resisted by friction (white arrows in Fig. 2d), Eq. (4). The sharp decrease in bond concentration at pocket margins is paralleled by a sharp increase in net membrane tension *σ* − *k*_*B*_*Tc* (purple map in Fig. 2b), which opposes the lateral expansion of the pocket footprint. Hence, mobile bonds control both the loss of cohesion leading to the nucleation of pockets when *T*_bonds_(*z*) ≈ *P*\*, and the lateral expansion of these pockets through *k*_*B*_*Tc*.

The nucleation and growth of the first row of pockets reduces hydraulic pressure locally, and hence this pocket front effectively constitutes a new edge of a smaller pristine adhesion patch (red map in Fig. 2b). The process then repeats with the nucleation and evolution of subsequent rows of pockets, Fig. 2b and Video 2, as long as the vesicle is sufficiently out of osmotic equilibrium to tear the adhesion apart. Our simulations exhibit profuse coalescence as well as events of pocket collapse, further discussed later.

### Principles of pattern selection

Varying various physical parameters such as *D, α*, μ, δΠ and *N*_*b*_, leads to a large diversity of fracture patterns, in terms of localization, size and spacing of pockets, and dynamics (SI). To systematically parse these behaviors, we identify the main non-dimensional numbers controlling the process.

Pocket formation requires that the osmotic shock is large enough to pressurize the interstice beyond the critical pressure of the effective adhesion potential *U*(*z*), hence defining the dimensionless osmotic shock 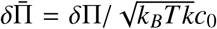, where the reference bond concentration is 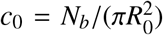 and *R*_0_ is the typical radius of the vesicle and the adhesion patch. By either changing the magnitude of the osmotic shock or the number of bonds, our simulations confirmed that 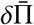 predicts the nucleation of pockets (SI).

Besides having a strong enough osmotic perturbation, pocket formation requires that this perturbation diffusively penetrates into the interstice fast enough compared to the time of overall vesicle osmotic relaxation by permeation τ_osm_ = *R*_0_/(*K*δΠ). This effect can be quantified by comparing *R*_0_ with the distance of diffusive osmolyte transport during τ_osm_ given by 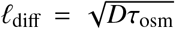, resulting in the dimensionless number 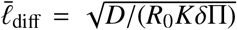. In agreement with this rationale, our simulations show that if this number is large, then pockets form throughout the adhesion patch. On the contrary, if 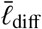 is small, pockets only form in a small region close to the edge or do not form at all, Fig. 3a. More quantitatively, we find a linear relation between 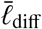 and the pocket penetration distance *d*_pen_ normalized by *R*_0_, Fig. 3a (inset).

**Figure 3:**
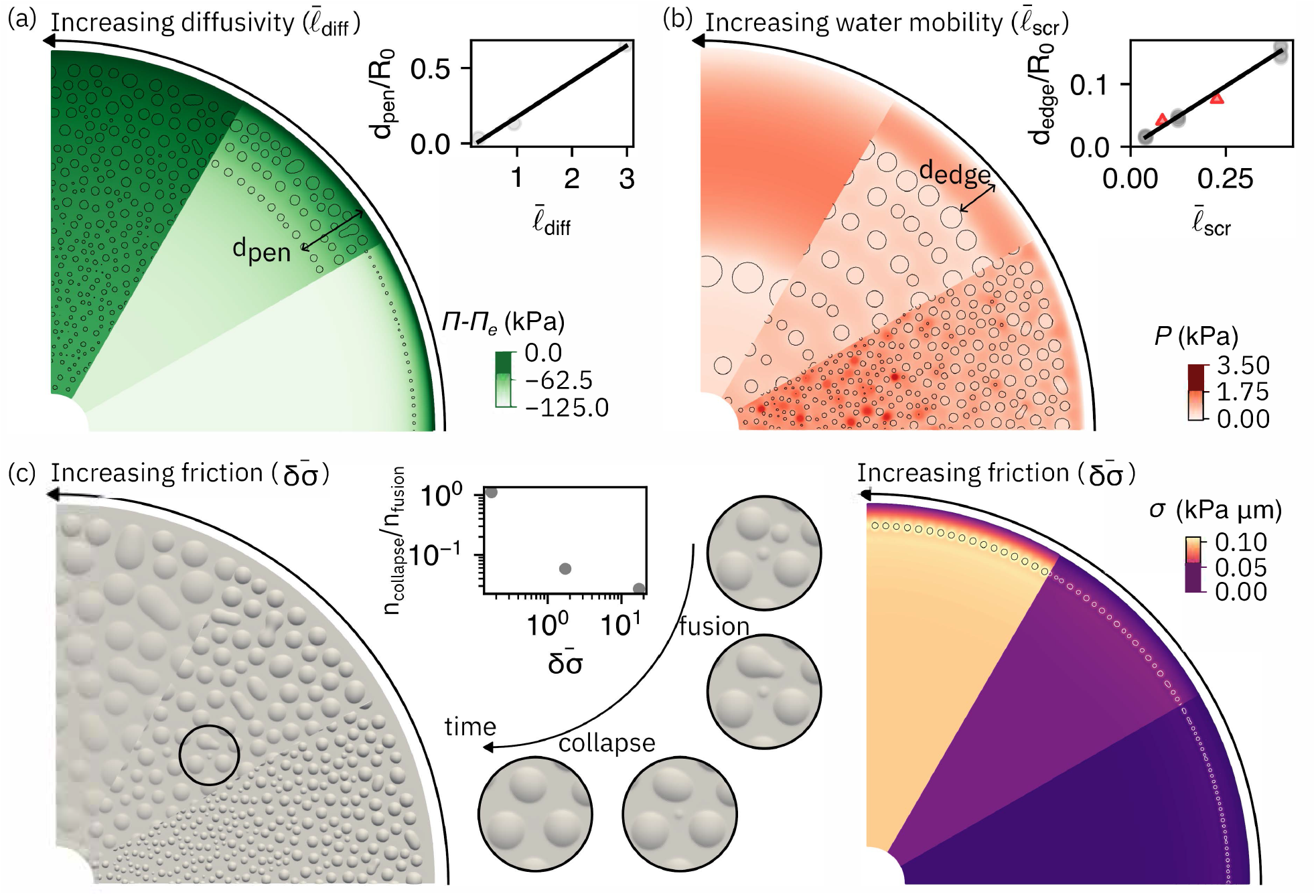
Pattern selection. (a) Osmotic pressure relative to the external medium for three different values of 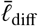 (0.30, 0.94, 2.98) obtained by changing *D*. Pocket boundaries are marked in black. The inset shows the penetration length *d*_pen_ (distance between the edge of the patch and the innermost pocket) normalized by patch radius as a function of 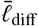. (b) Hydraulic pressure relative to external medium for three different values of 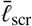 (0.04, 0.12, 0.39). The inset shows the distance between the outermost pocket and the edge *d*_edge_ as a function of 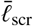. Circles mark simulations where *α* is changed and triangles where *K* is changed to modify 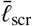. (c) Top view of 3D shape of the pockets for three different values of 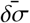 (0.17, 1.73, 17.32) obtained by changing μ. The inset shows the relative number of collapses and fusions during pattern coarsening as a function of 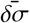. Zoom plots show sequence where both fusion and collapse take place. The map on the right shows bare membrane tension right after nucleation, showing higher friction generates larger tension gradients.

The condition that 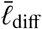 is large enough guarantees significant water permeation into the interstice, and hence is required for pocket formation but not sufficient. For this efflux to pressurize the interstice, it should be opposed by hydraulic resistance, which can be quantified by the dimensionless hydraulic screening length 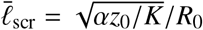^34^. At a distance smaller than ℓ_scr_ from the adhesion edge, water can easily leave the interstice by Darcy flow. Thus, if 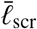 is comparable or larger than 1, we expect very low hydraulic confinement and no pocket formation. For smaller values, we expect ℓ_scr_ to determine the distance between the first row of pockets and the edge, as well as the separation between subsequent rows of pockets. Furthermore, since the dynamics of pocket growth requires water volume reconfigurations by permeation and Darcy water flows, Fig. 2c and Video 3, we expect ℓ_scr_ to dictate the typical size of pockets. These arguments are confirmed by our simulations, including a quantitative linear relation between the normalized distance to the edge of the first pockets *d*_edge_/*R*_0_ and 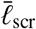, Fig. 3b.

Finally, lipid membranes being nearly inextensible, pocket formation requires recruitment of membrane area from the free-standing part of the vesicle and pocket reorganization from nearby regions, Fig. 2d. When the dimensionless hydrodynamic length 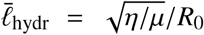 is ≪ 1 (∼ 10^−4^ in our system), the dominant mechanism dragging membrane flow is friction, which hinders membrane tension equilibration^35^. To estimate induced differences in membrane tension δ*σ*, we note that tension gradients first develop between the initial row of pockets and the edge of the adhesion patch, and hence have a characteristic buildup time of 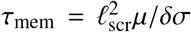. On the other hand, we estimate the time for pocket growth as τ_growth_ = ℓ_scr_/*v*_*p*_ = ℓ_scr_/(*K*δΠ). Equating these two times and non-dimensionalizing by the osmotic tension scale, we find the dimensionless quantity 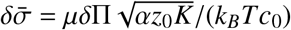 characterizing the frictional opposition to pocket growth. In agreement with these arguments, our simulations show that for small 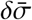, gradients in *σ* are small, and since *γ* = *k*_*B*_*Tc* is nearly uniform, all pockets exhibit similar contact angles (Fig. 3c) supplementary to *θ* (Fig. 2a) according to the Young-Dupré relation. In contrast, for large 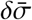, strong tension gradients develop increasing towards the interior of the patch. Tension being large, the system develops very large and shallow pockets that enclose water volume resulting from vesicle efflux with very small excess area^17^ (Fig. 3c). Furthermore, the gradient of tension is mirrored by a gradient in contact angles. The contact angles in our experiments are large, and hence we infer that our system operates in a low friction regime.

Regarding the coarsening mechanism, our simulations exhibit coexistence of pocket fusion and collapse (Fig. 3c). For low 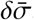, the number of collapse events is similar to the number of fusion events, whereas large 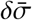 favors fusion as pockets grow by laterally expanding their footprint area. Compared to experiments, a significant difference is that in our simulations nearby pockets readily fuse, whereas observations of stable pairs of pockets at very close distance are common (Fig. 1c,d). We attribute the barrier to pocket fusion to the presence of trapped adhesion molecules between pockets.

In summary, the dimensionless numbers 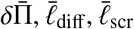 and 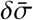 delineate non-generic conditions for pocket formation, imposing conflicting requirements on various parameters such as size, *K*, δΠ or *c*_0_. For instance, increasing *K* by embedding aquaporins in the membrane^23^ reduces both ℓ_diff_ and ℓ_scr_, while pocket formation requires the first of these lengthscales to be large and the second to be small compared to *R*_0_. These dimensionless numbers also control the nature and dynamics of the pattern of hydraulic cracks, providing a means to estimate the poorly characterized transport parameters (diffusivity, water mobility and membrane friction) in cell-cell or artificial membrane adhesion clefts^34^.

### Experimental control of fracture patterns

To test the theoretical predictions, we modify experimentally accessible parameters. We start by changing the strength of the osmotic shock. As predicted by theory and simulations, we observe nearly no pockets for the smallest shock, δΠ = 25 mM, whereas detached area increases with increasing δΠ (Fig. 4a). However, this increase of detached area does not take place throughout the patch and remains largely confined to the periphery, consistent with the fact that 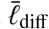 decreases with increasing δΠ.

**Figure 4:**
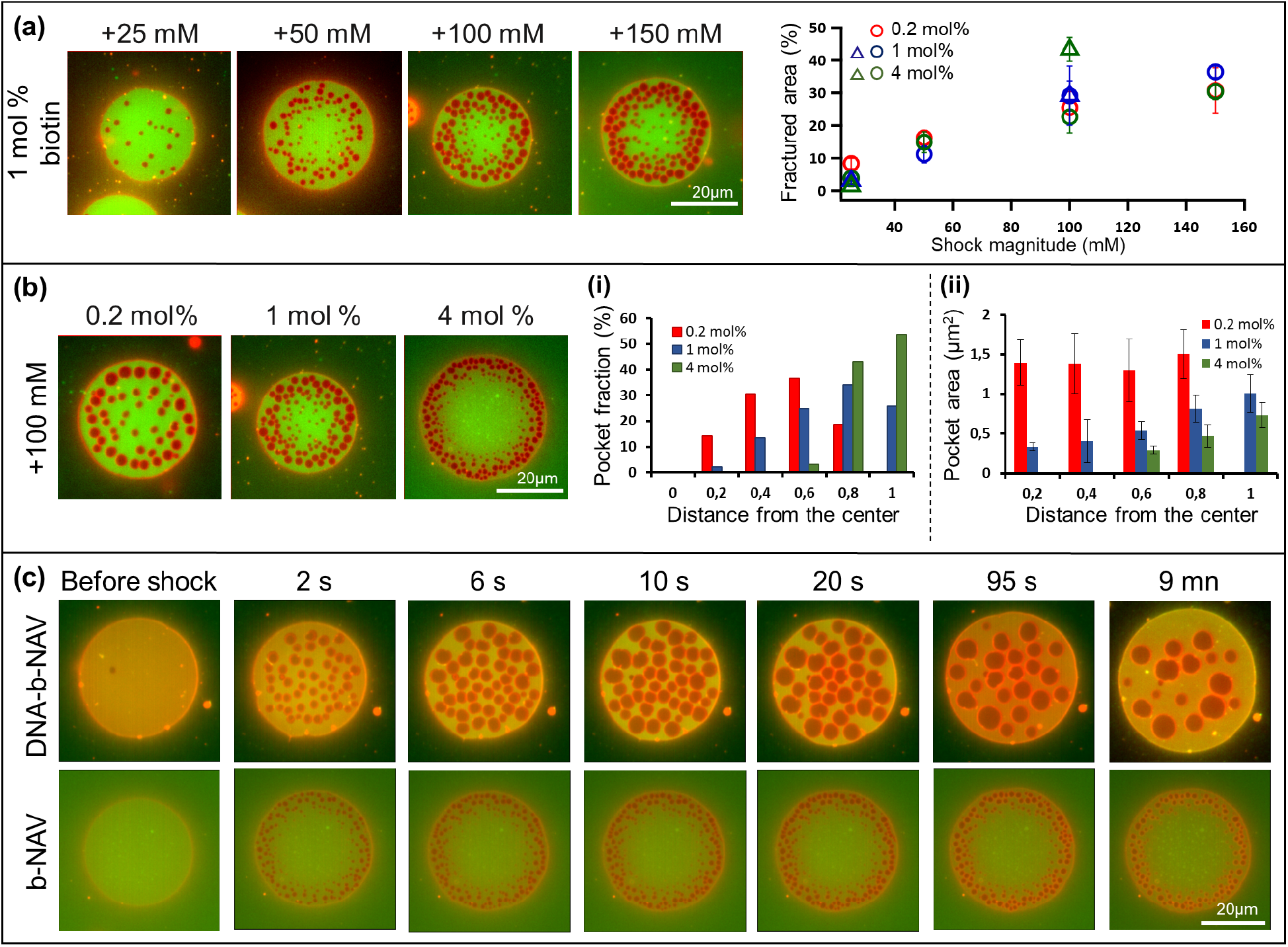
Hydraulic fracture patterns for various experimental parameters, quantified when all pockets have appeared and before they start coarsening. (a) Varying osmotic pressure. Left: Epi-fluorescent images of the GUV-SLB adhesion zone for 1 mol% biotinylated lipids at various shocks. Right: Total fractured area as a function of shock magnitude. Round circles and triangles mark b-NAV and DNA-b-NAV bonds, respectively. (b) Varying bond density. Left: Epi-fluorescent images of adhered vesicles with 0.5, 1 and 4 mol% biotinylated lipids subject to 100 mM hyper-osmotic shock. (i) Number fraction of pockets and (ii) average pocket size as a function of distance from the patch center, quantified in 0.2 radius fraction intervals. (c) Time lapse epi-fluorescent images of adhered vesicles (4 mol%) with and without DNA bond spacers before and after the application of 100 mM osmotic shock. Membrane appears in red and the NAV bonds in green.

We then change the bond concentration. A larger bond concentration increases molecular crowding in the thin interstitial space, and hence should impair transport and reduce diffusivity *D*^36^ and Darcy mobility *α*^37^. Consistent with reduced diffusivity, and hence smaller 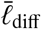, pockets form in an increasingly narrow peripheral region as bond concentration increases (Fig. 4b-i). Also consistent with reduced Darcy mobility, and hence smaller 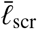, pockets appear smaller, closer together and closer to the edge of the patch (Fig. 4b-ii).

To examine the opposite regime of reduced molecular crowding, and hence of large 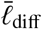 and 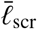, we note that in b-NAV bonds the bulkiest molecule is the NAV protein. Therefore, intercalation of thin DNA linkers between lipids and biotins not only increases membrane separation (from 5.6 to 25.6 nm) but also reduces crowding. Consistent with this rationale, in adhesions with DNA-b-NAV bonds, pockets form uniformly throughout the adhesion patch and are much larger, in sharp contrast with the fracture pattern of an otherwise similar adhesion with b-NAV bonds (Fig. 4c).

### Pocket dynamics at GUV-GUV interfaces and at long times

In the following we discuss the evolution of pockets over longer times, and compare pockets formed between vesicles and supported lipid bilayers (GUV-SLB) as well as between two vesicles (GUV-GUV). Following pattern formation, pockets on the GUV-SLB adhesion patch appear immobile for all bond densities and do not change significantly their shape and size over a period of at least 6-7 min. Occasionally, we observe coalescence between adjacent pockets and pockets collapse, especially for smaller pockets and those near the adhesion rim (Fig. 1 and SI). In systems with high water mobility, as in the case of the longer DNA-b bonds, pockets remain immobile but they can discharge in the outer medium and collapse through water diffusion (Video 4). Pockets at the GUV-GUV adhesion patch on the other hand are highly mobile and exhibit significant Brownian motion (Fig. 5a-i, Video 5). Over a comparable time period they coalesce into a single large pocket, which can discharge its content and sometimes bud upon contact with the external rim of the adhesion patch (Fig. 5a-ii). Such enhanced mobility and coalescence are favoured for smaller bond density and at lower osmotic shocks. Larger osmotic shocks lead to the formation of a packed pattern of pockets with limited space for their diffusion (Video 6). Our FRAP experiments show that the mobility of lipids and NAV bonds is greatly reduced in supported vs unsupported membranes (SI), and hence we attribute impaired pocket mobility in GUV-SLB adhesions to substrate drag. For the same reasons, lipid domains in ternary membrane mixtures are mobile in vesicles and immobile in SLBs^38^. Mobility of lipids and NAV further decreases with increasing bond density^21^ (SI).

**Figure 5:**
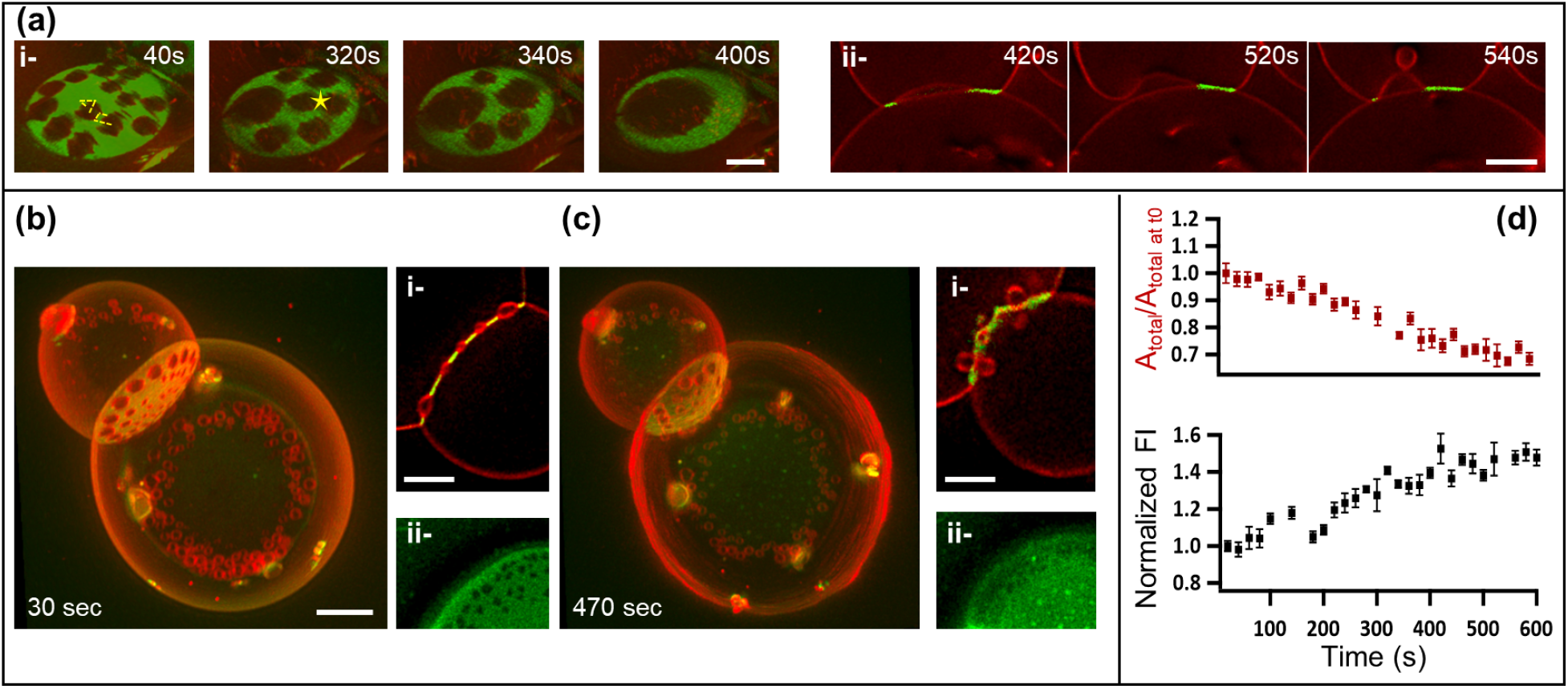
Long term dynamics of pockets at the vesicle-vesicle adhesion patch. (a) Coalescence of water pockets in the adhesion zone between two vesicles linked at 0.5 mol% biotinylated lipids and subject to 50 mOsm osmotic shock, i-3D reconstitution of the adhesion zone, ii-cross-section images of the same adhesion zone some time later. The dashed lines show an average pocket displacement of 2 μm within several 20 sec intervals. The star indicates coalescence of pockets. (b,c) Budding of water pockets between two vesicles linked at 4 mol% biotinylated lipids and subjected to 100 mOsm osmotic shock at 30 (b) and 470 (c) sec after the shock. The insets show (i) a cross-section of the GUV-GUV adhesion zone and (ii) of the GUV-SLB adhesion zone. Scale bar in all images is 4 μm. (d) Plots of the total GUV-GUV adhesion area and of the mean NAV fluorescent intensity in the adhesion zone, corrected for photo-bleaching, sampled at 10 different locations. The values in both plots are normalised to the initial values, where t=0 is the time at which imaging starts (approximately 30 sec after the osmotic shock).

In addition to the coarsening through coalescence and collapse, we often observe pocket budding, both from GUV-GUV and GUV-SLB adhesion patches (Fig. 5b,c and Video 7). Buds remain to hover in the vicinity of the adhesion patch, suggesting that they are still connected to it, but are not reabsorbed within the observation time. To understand the mechanism leading to closing of pockets into buds, we first examine whether the lateral osmotic pressure of bonds may push neck edges to reduce bond density. Our observations of GUV-GUV interfaces show that budding is accompanied by significant shrinking of the adhesion patch area and by an increase in bond density (Fig. 5d, Video 7), ruling out this possibility. We then note that transitions from shallow pockets to buds of comparable volume in an adhered membrane are generically caused by an increase in excess membrane area or a reduction of membrane tension^17^. Since tension in an adhered vesicle is determined by the strength of adhesion, we examine the GUV-SLB adhesion patch where bonds are less mobile. At long times, we often observe membrane unbinding by bond breaking, detected by a loss of NAV signal, as well as an accumulation of NAV bonds at the location of the buds (Fig. 5c (ii), and SI). Furthermore, the process of pocket budding, shrinking of the GUV-GUV patch and loss of NAV signal at the GUV-SLB interface, coincide with strong vesicle fluctuations (Fig. 5c).

These observations show that our system transits from a strong adhesion regime with intact bonds and reduced fluctuations before the shock and at the initial stages of pattern formation, to a weakened adhesion regime with bond breaking at the GUV-SLB interface and reduced tension, leading to prominent fluctuations and budding at GUV-GUV interfaces. This phenomenology is reminiscent of earlier studies on the behaviour of bonds under force-induced membrane detachment, where mobile bonds are displaced and concentrate in the shrinking adhesion patch, and immobile or less mobile bonds tend to break under a pulling forces^24;33;39^. Furthermore, previous studies have shown that membrane fluctuations give rise to an entropic repulsion force, which can modify the density of bonds and their effective binding strength^24;40^, and can even trigger membrane unbinding in the case of low adhesion strength^41;42^. We hypothesize that the reduction of membrane tension following pocket formation enables initially small fluctuations, which over a long period of time can break a certain amount of bonds at the edge of the tight adhesion zone. Because the length of this edge is increased by the presence of pockets, this mechanism is enhanced. Bond breaking then reduces adhesion tension, leading to further reduction of membrane tension, enhanced fluctuations, thereby establishing a positive feedback loop. Our experiments thus demonstrate a novel transition from strong to weak adhesion mediated by hydraulic fractures (in the GUV-SLB interface) and a mechanism of irreversible budding of hydraulic pockets driven by tension reduction (in the GUV-GUV and GUV-SLB interfaces).

## Conclusions

In this work, we study the hydraulic fracturing of lipid vesicles strongly adhered through mobile bonds upon osmotic deflation. The resulting patterns of water pockets and their evolution resemble the process of microlumen formation and coarsening in embryonic tissues^8^. By combining theoretical modeling, numerical simulations and experiments, we identify the physical principles controlling nucleation, spatial pattern and dynamics of hydraulic cracks. The conditions for pattern formation are non-generic, and require an intermediate degree of confinement of the adhesion cleft. If too confined, osmotic imbalances cannot penetrate the interstice, whereas if insufficiently confined, water efflux can escape the cleft without compromising adhesion. We further show that over time, the presence of pockets can weaken the adhesion patch, lower membrane tension, and lead to budding of pockets, akin to precursors of endocytic vesicles in our minimal in vitro system.

Our work provides a physical basis for reconfigurations of cell-cell adhesions. In general, biological patterning and reshaping during development results from an interplay between mechanics and biochemical regulation^43^. In the context of luminogenesis, our work identifies the physical rules enabling the initial patterning of profuse hydraulic cracks at every cell-cell junction, on top of which the previously identified mechanism guiding coarsening by gradients of cell surface tension can act to position the blastocoel^8^. For instance, our results suggest that rather than hydraulic confinement by tight junctions at the cell-medium interface, profuse cracking requires reduced water mobility throughout cell-cell adhesions in the embryo, and that irreversible budding of pockets is avoided by sufficiently large cellular tension. In the context of adhesion remodeling, our work suggests that such irreversible budding triggered by reduced membrane tension may constitute a physical pre-patterning mechanism for endocytic vesicles, subsequently tamed by known biochemical regulatory pathways^44;45^.

## Methods

### Consumables

1,2-dioleoyl-sn-glycero-3-phosphocholine (DOPC), 1,2-dioleoyl-sn-glycero-3-phosphoethanolamine-N-(cap biotinyl) (sodium salt) (b-DOPE) and 1,2-dipalmitoyl-sn-glycero-3-phosphoethanolamine-N-(lissamine rhodamine B sulfonyl) (ammonium salt) (Rhod-DPPE) were all purchased from Avanti Polar Lipids (Alabaster, AL) and used without further purification. NeutrAvidin Protein, DyLight 488 (NAV) was purchased from Thermo Fisher Scientific. Chloroform, trizma hydrichloride (Tris·HCl), glucose and sucrose were purchased from Sigma Aldrich. Microscope slides and cover glasses from VWR (catalog no. 48366 045) were used. For the preparation procedure of GUVs, we used Indium Tin Oxide coated glasses (ITO glasses) from Delta Technologies (no. X180). Double stranded DNA carrying a biotin moiety on one side and a double cholesterol anchor on the other side were gift from L.Di Michele lab (Department of Chemical Engineering and Biotechnology, University of Cambridge, UK). The biotin moiety allows the DNA to bind to the Neutravidin protein and the cholesterol functionalization renders the nanostructures amphiphilic and able to spontaneously insert into the hydrophobic core of the lipid bilayer membrane.

### Substrate preparation and chamber

Glass cover slides were washed several times with isopropanol and ultrapure water (18.2 MΩ, 0.5 ppm organics, Merck Millipore), dried with nitrogen flow and were then further cleaned by exposing to low-pressure air plasma, at a pressure of 1mbar and power of 300 watts (VacuLAB Plasma Treater, Tantec), for 40 sec to render the cover-slide clean and hydrophilic. The experimental chamber was assembled by sticking a PDMS spacer onto the cleaned glass substrate. The total volume of the chamber was 500 μl.

### Preparation of supported lipid bilayer (SLB)

SLB were prepared using vesicle fusion technique as described previously^46;47;48^. Briefly, a thin film of 2mg lipids formed by 99.5; 98.7 or 95.7 mol% DOPC, 0.2, 1 or 4 mol% b-DOPE, respectively and 0.3 mol% Rhod-DPPE were dried in a vacuum dessicator overnight on the wall of glass vial. The following day, the dried lipid film is rehydrated in lipid buffer (10mM Trizma base; 150 mM NaCl and 2mM CaCl2, pH ≈ 7.5) to a final concentration 1mg/mL. The resulting suspension is then sonicated using a tip sonicator operated in a pulsed mode at 20% power for 10 min with refrigeration to generate small unilamellar vesicles (SUV’s) from the lipids. The solution is then centrifuged at 1000 rpm for 10 mn in an Eppendorf centrifuge to remove titanium particles. SUV suspensions were stored at 4°C under nitrogen and used within a week. A dilution of the SUVs suspension with lipid buffer at a 1 : 4 volume ratio is spread over the clean hydrophilic glass cover-slide in a final 200 μl volume created by the PDMS chamber (see above). Incubation for about 30-60 min results in the formation of a supported lipid bilayer. The SLB was then thoroughly washed with glucose solution having a concentration of 300 mM (isotonic relatively to the sucrose solution in which the GUVs have been prepared). This is done to remove unfused SUV’s and the lipid buffer.

### Preparation of giant unilamellar vesicles (GUV)

GUVs owing the same composition as that of the SLB were produced via electroswelling^49;50;51^. Briefly, 50 μl of the solution containing the lipid mixture were dispersed on two titanium oxide-coated glass slides. The lipid coated slides were dried in a vacuum desiccator for at least 5 hours to ensure chloroform evaporation. The dried coated lipid slides are put together with a Teflon spacer to form a capacitor cell. The conductive side of the two slides were faced inward and fixed with a clamp to form a chamber. The chamber was then filled with 300 mM sucrose solution, and an alternating current of 10 Hz and 2V peak to peak amplitude was applied across the chamber and kept overnight. The GUV’s were then extracted from the chamber, stored in an Eppendorf vial and used within 2-3 days.

### Immobilization of giant unilamellar vesicles

To bind GUVs to SLB with biotin-Neutravidin bonds (b-NAV), the GUVs and SLB were prepared from the same lipid stock solutions of DOPC and b-DOPE as described above. Before vesicle adhesion, the SLB was incubated with an excess of NAV at a final concentration of 60 μg/ml for 30 mn and then rinsed with glucose 300 mM solution to remove excess protein. Following that, 2-5 μl of GUV solution was added to the chamber and incubated for 30 mn to allow the GUVs to sediment on the SLB and form adhesion site with each other. The solution is then washed carefully with 300 mM glucose solution to remove unbound vesicles.

To bind GUVs to SLB with biotynilated DNA-Neutravidin bonds (DNA-b-NAV), we followed the experimental procedure described in Amjad *et al*.^52^. The DNA constructs were stored in DNA buffer at a concentration of 5 μM. SLBs and GUVs were prepared from a lipid mixture of 99.7mol% DOPC and 0.3% mol Rhod-DPPE as above. The SLBs were rinsed with DNA buffer (300 mM) (Tris EDTA (1X); 100 mM NaCl and 87 mM glucose) instead of glucose (300mM). To achieve respectively 1, 4, 6 or 8 mol% b-DNA linker density in both SLB and GUVs; 0.234, 1, 1.5 or 2 μl of the DNA constructs solution was added together with 0.5 μl of GUV solution to the SLB. The chamber was incubated for 1 hour to allow grafting of the DNA to the lipids. A desired amount of the NAV solution was then added to the chamber to achieve a ratio of NAV/DNA-b of 1/4, and was left incubating for 1hour. This in theory allows all NAVs to bind 4 DNA-b constructs. The solution is then washed carefully three times with 300 mM DNA buffer solution.

### Osmotic shocks

By the time GUVs were bound to the SLB, all samples had a final volume of 400 μL. To subject the b-NAV GUVs to hyperosmotic shocks of 25, 50, 100 and 150 mM osmotic shocks, half of the volume of the chamber (200 μL of the 300mM osmolarity) was replaced by glucose solutions of 350, 400, 500 and 600 mM, respectively. For the DNA-b-NAV GUVs, the shock solutions were 350 mM (Tris EDTA (1X) + 100 mM NaCl + 137 mM glucose), 400 mM (Tris EDTA (1X) + 100 mM NaCl + 187 mM glucose); 500 mM (Tris EDTA (1X) + 100 mM NaCl + 287 mM glucose) and 600 mM (Tris EDTA (1X) + 100 mM NaCl + 387 mM glucose), respectively.

The precise osmolarity of the shock solutions was measured for each experiment with an osmometer (Osmomat 3000, Gonotec GmbH, Berlin, Germany). After the addition of the shock solution, the chamber was covered to prevent further osmolarity changes due to evaporation.

### Imaging and analysis

The imaging of the adhesion zone between the SLB and GUV throughout the osmotic shock was performed with an inverted optical microscope Nikon Eclipse Ti-E and a 60x numerical aperture, oil immersion objective in combination with an Andor camera Neo 5.5 sCMOS (Oxford Instruments). The integrated perfect focusing system (PFS) in the microscope allows us to follow automatically the surface which change its focal plane during the application of the osmotic shock. The open source image processing package FIJI was used for the image analysis. The changes in the adhesion area and intensity in response to the osmotic shock are performed by first subtracting the background of the fluorescence images and then applying an appropriate thresholding to generate a binary stack. Analyze particle function was then used to obtain the total and intimate adhesion area and intensity. The parameters reported in the paper are averages calculated from at least 2 samples, with at least 3 vesicles each. Confocal images were acquired using a Zeiss LSM 880 Fast AiryScan and a Plan-Apochromat 63x numerical aperture 1.4 Oil immersion objective. The three-dimensional (3D) reconstruction using the confocal stack was done using a Fiji plugin (ClearVolume)^53^. We always closed the chambers during imaging of the vesicles in order to avoid convection and large-scale drifts.

## Simulations

The equations describing the time-evolution of the system involve a set of fields (*z*, ***v***, *P*, Π, *σ*) in the patch 𝒟(*t*) and the variables (*R, θ, P*_*i*_, Π_*i*_, *σ*_*v*_) representing the state of the vesicle (see Box 1). We integrate these equations in time in a staggered way, by first solving the equations for (*z*, ***v***, *P*, Π, *σ*) with a backward Euler approximation assuming fixed values of (*R, θ, P*_*i*_, Π_*i*_, *σ*_*v*_) and then solving for the vesicle variables assuming fixed values for (*z*, ***v***, *P*, Π, *σ*). To discretize (*z*, ***v***, *P*, Π, *σ*) in 𝒟(*t*) we consider a triangular mesh and use a second-order Lagrangian interpolation for (*z*, ***V***, *P*, Π) and a first-order Lagrangian interpolation for *σ* where here ***V*** = ***v*** + *v*_*n*_***N*** is the three-dimensional velocity of lipids. To recover *z* from ***V***, we note that since ∂_*t*_*z* = *v*_*n*_, we can approximate *z*(*t* + Δ*t*) ≈ *z*(*t*) + (***V*** · ***N***) Δ*t*. To compute the tangential velocity ***v***, we project ***V*** onto Γ_*t*_. To solve the balance of forces on the membrane we follow the procedure detailed in^32^. The equations are then solved using a finite element method with the boundary conditions discussed in Box 1 and implemented in hiperlife^54^.

## Supporting information

Pocket formation and coarsening between a vesicle and a SLB, both containing 0.2 mol% biotinylated lipids and subjected to 100 mM hyper-osmotic shock.

Simulation of pocket formation and evolution, corresponding to Fig. 2b.

Water permeation and interstitial flow during pocket formation and evolution, corresponding to Fig. 2c.

Pocket dynamics between a vesicle and a SLB, containing 6 mol% biotinylated DNA and subjected to 100 mM hyper-osmotic shock.

Mobility of pockets between two adhered vesicles, both containing 0.5 mol% biotinylated lipids and subject to 50 mM hyper-osmotic shock.

Crowding and restricted mobility of pockets between two adhered vesicles, both containing 0.5 mol% biotinylated lipids and subject to 100 mM shock.

Pocket budding and shrinking of the adhesion zone between two adhered vesicles, both containing 0.5 mol% biotinylated lipids, at later stages.

---

**Supplementary Table 1:**
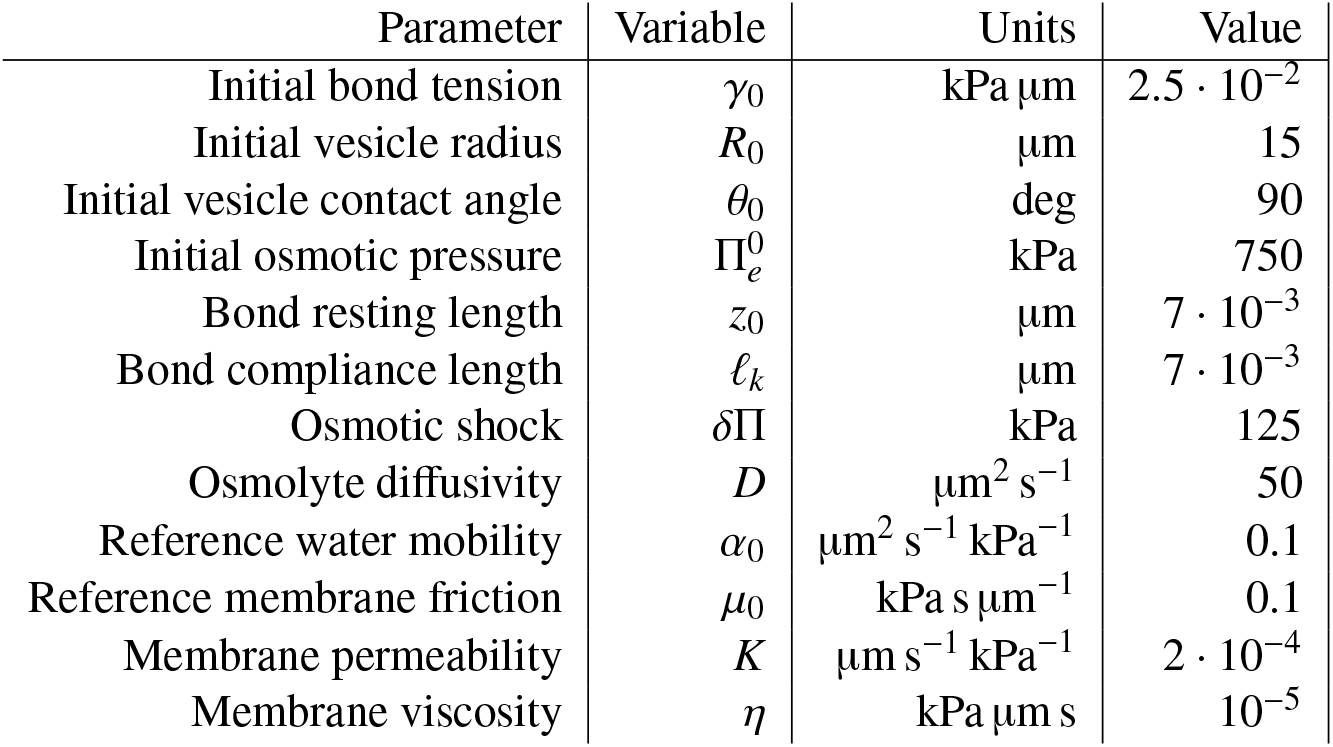
Table of parameters used for the simulation in Fig. 2 and Video 2. The data characterizing the initial state prior to the shock (first three lines) can be easily mapped to the initial data given in Box 1 (number of bonds *N*_*b*_, number of trapped osmolytes *N*_*o*_, vesicle surface area *S*). The bond compliance length measures is the amplitude of typical bond thermal fluctuations and also the critical separation distance for loss of stability of an adhesion. It is given by 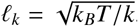. The osmotic shock δΠ = 125 kPa corresponds to 50 mM.

## Video captions

**Video 1:** Pocket formation and coarsening between a vesicle and a SLB, both containing 0.2 mol% biotinylated lipids and subjected to 100 mM hyper-osmotic shock, corresponding to Fig. 1c-d.

**Video 2:** Simulation of pocket formation and evolution, corresponding to Fig. 2b.

**Video 3:** Water permeation and interstitial flow during pocket formation and evolution, corresponding to Fig. 2c.

**Video 4:** Pocket dynamics between a vesicle and a SLB, containing 6 mol% biotinylated DNA and subjected to 100 mM hyper-osmotic shock.

**Video 5:** Mobility of pockets between two adhered vesicles, both containing 0.5 mol% biotinylated lipids and subject to 50 mM hyper-osmotic shock, corresponding to Fig. 5a.

**Video 6:** Crowding and restricted mobility of pockets between two adhered vesicles, both containing 0.5 mol% biotinylated lipids and subject to 100 mM hyper-osmotic shock.

**Video 7:** Pocket budding and shrinking of the adhesion zone between two adhered vesicles, both containing 0.5 mol% biotinylated lipids, at later stages following a 100 mM hyper-osmotic shock.

## Notes

### Competing Interest Statement

The authors have declared no competing interest.

